# Latent developmental and evolutionary shapes embedded within the grapevine leaf

**DOI:** 10.1101/018291

**Authors:** Daniel H. Chitwood, Laura L. Klein, Regan O’Hanlon, Steven Chacko, Matthew Greg, Cassandra Kitchen, Allison J. Miller, Jason P. Londo

## Abstract

- Across plants, leaves exhibit profound diversity in shape. As a single leaf expands, its shape is in constant flux. Plants may also produce leaves with different shapes at successive nodes. Additionally, leaf shape varies among individuals, populations, and species due to evolutionary processes and environmental influences.
- Because leaf shape can vary in many different ways, theoretically the effects of distinct developmental and evolutionary processes are separable, even within the shape of a single leaf. Here, we measure the shapes of >3,200 leaves representing >270 vines from wild relatives of domesticated grape *(Vitis* spp.) to determine if leaf shapes attributable to genetics and development are separable from each other.
- We isolate latent shapes (multivariate signatures that vary independently from each other) embedded within the overall shape of leaves. These latent shapes can predict developmental stages independent from species identity and vice versa. Shapes predictive of development are then used to stage leaves from 1,200 varieties of domesticated grape *(Vitis vinifera),* revealing that changes in timing underlie leaf shape diversity.
- Our results indicate distinct latent shapes combine to produce a composite morphology in leaves, and that developmental and evolutionary contributions to shape vary independently from each other.

## Introduction

Leaf morphology represents a beautiful and tangible example of the infinite phenotypic possibilities in nature. Underlying leaf shape diversity is a quantitative genetic (Langlade *et al.*, 2005; Kimura *et al.*, 2008; Tian *et al.*, 2011; Chitwood *et al.*, 2013) and developmental genetic (Bharathan *et al.*, 2002; Kim *et al.*, 2003; Blein *et al.*, 2008) framework. It is possible that aspects of leaf shape are functionally neutral and reflect developmental constraint (Chitwood *et al.*, 2012a; 2012b), but numerous hypotheses about the function of different leaf shapes exist, including how shape impacts thermal regulation, hydraulic constraints, light interception, biomechanics, and herbivory (Parkhurst and Loucks, 1972; Nicotra *et al.*, 2011; Ogburn and Edwards, 2013). Fossil leaf size and dissection are correlated with the paleoclimate (Bailey and Sinnott, 1915; Wolfe, 1971; Greenwood, 1992; Wilf *et al.*, 1998), a relationship that persists in extant taxa (Peppe *et al.*, 2011), and with implications for the chemical, structural, and physiological economics of leaves (Wright *et al.*, 2004). Correspondingly, functional traits related to leaf shape display phylogenetic signal in some clades (Cornwell *et al.*, 2014). Understanding the spatial and temporal patterns of leaf shape variation is a central theme in studies focusing on plant biodiversity, impacts of global climate change, and agricultural efficiency.

Leaf shape varies not only across evolutionary timescales and within a functional ecological context, but during development as well. Two distinct temporal processes regulate leaf shape during development. 1) The shape of individual leaves is in constant flux as local regions within the leaf expand at different rates. This phenomenon, allometric expansion, was explored as early as Hales’ *Vegetable Staticks* (1727). Using a grid of pins, regularly spaced puncture points in fig leaves were tracked to determine if their relative spacing changed during development. The same experiment can be microscopically studied using fluorescent particles today (Remmler and Rolland-Lagan, 2012; Rolland-Lagan *et al.*, 2014). 2) The leaves that emerge at successive nodes differ in their shape, as the shoot apical meristem from which they derive transitions from a juvenile to adult stage of development. This process, heteroblasty, can affect other features of leaves besides shape, such as cuticle and trichome patterning (Goebel, 1900; Ashby, 1948; Poethig, 1990; Kerstetter and Poethig, 1998; Poethig, 2010).

The developmental stage of a leaf and the position of the node from which it arises (leaf number) are distinct temporal factors affecting leaf shape. Genetic changes in the timing of either process (e.g., protracted development of individual leaves or precociously adult leaf morphology in early nodes) between species can lead to evolutionary differences in leaf shape, a process known as heterochrony (Cartolano et al., 2015). The timing of these processes can be changed non-genetically as well, through responses to environmental changes during the lifetime of a plant, known as plasticity (Allsopp, 1954; Diggle, 2002). Distinguishing effects of developmental stage from leaf number provides mechanistic insights into how genetic changes during evolution or plastic changes in response to the environment are achieved (Jones, 1993; 1995). For example, recent work has linked molecular pathways regulating the timing of shape changes throughout the shoot (miR156/172 and their targets) with leaf morphology (CUC-induced serrations) through a mediator (TCPs) (Rubio-Somoza *et al.*, 2014).

If distinct molecular pathways affect different traits within the infinite features defining the architecture of a leaf, then the outline and venation topology of leaves should theoretically be decomposable into latent (hidden) shapes; i.e., multivariate morphological signatures that vary independently from each other (Chitwood and Topp, 2015). For example, shape differences defining species, as well as shape differences defining developmental stage, may be detectable within each leaf and subsequently isolated from each other. Shape differences that define species, regardless of developmental context, may vary distinctly from those features that define developmental context, regardless of species. These shapes are latent because, while they are present in each leaf, they manifest within the context of other factors that affect shape as well. Latent shapes globally affect leaf shape in different ways, and each of the features comprising the latent shapes alone do not necessarily discriminate the effects of genetics or development. From this perspective, the shape of a given leaf—from any species, from any time point during development or any position in the plant—would result from the confluence of latent shapes regulated by these processes. The single organ that we call a leaf would actually be a composite of latent features that vary by genetic versus developmental effects, independently from each other.

Grapevine (*Vitis* spp.) leaves exhibit a breathtaking range of variation in leaf shape, making this genus ideally suited to explore latent shapes resulting from evolutionary and developmental processes **(Fig. 1)**. Taxonomists studying *Vitis* have used variation in leaf lobing and leaf margins to delimit the nearly 60 species in the genus (Moore, 1991; Ren and Wen, 2007). Leaf shape is also important in assessing intraspecific variation in the European grapevine (*Vitis vinifera* subsp. *vinifera),* which is grown around the world for wine-making and table grapes. Unique among crops, variation in grape leaf shape (together with other vine features) is used by viticulturists to quantitatively classify grape varieties, a field known as ampelography (αμπελος, “vine” and γραφος, ‘writing’) (Galet, 1952). Grape leaves have several homologous points amenable to landmark-based analyses (Galet, 1979; Chitwood *et al.*, 2014), increasing the biological interpretation of morphometric data compared to species with stochastic venation topologies or limited homology (such as *Arabidopsis,* tomato, *Antirrhinum,* etc.). Like Hales’ grid of pins (1727), naturally homologous points in grape leaves allow developmental stage and leaf number effects to be quantitatively tracked, potentially revealing separable latent processes contributing to the composite morphology known as a leaf.

**Figure 1:**
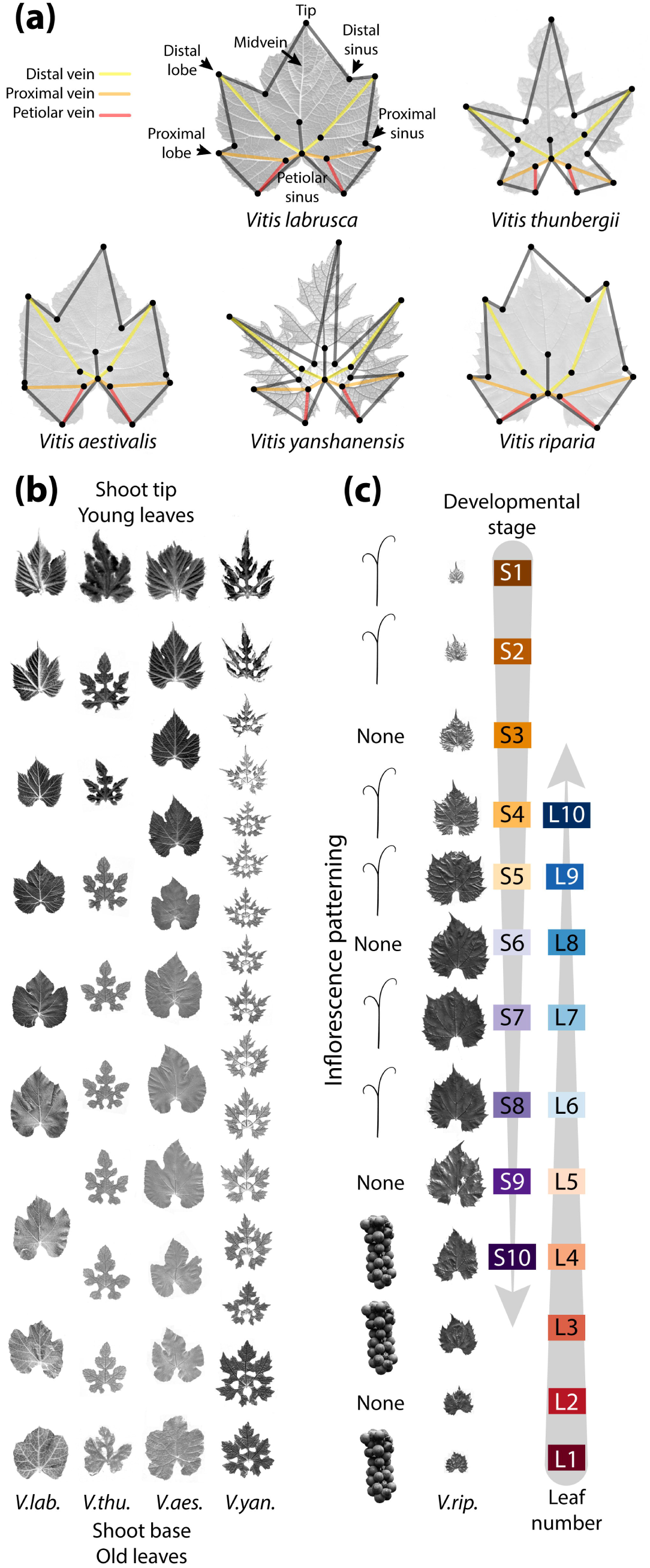
Morphological features defining the temporal development of *Vitis* leaves. (a) All *Vitis* leaves possess distal (yellow), proximal (orange), and petiolar (red) veins, as well as distal and proximal lobes and sinuses and a petiolar sinus. 17 homologous landmarks (black circles) were used in this study. Morphologically diverse species are shown, (b) Leaf series, scaled by the length between distal lobe tips, showing developmental stage and leaf number shape variance, (c) Unsealed *V. riparia* leaves showing changes in size at the shoot tip and base. Developmental stage (Sn) is measured starting from the shoot tip and leaf number is measured from the shoot base (Ln). Inflorescences, opposite leaves, skipping every third node, transform from clusters into tendrils from the shoot base to tip.

## Methods and Methods

### Germplasm, sample collection, and scanning

Over 270 vines in the USDA *Vitis* germplasm collection in Geneva, NY were sampled in June 2013. Using an automatic label maker, vine identification number and the identity of organs (or lack thereof) at each node, beginning with the first sampled leaf at the tip, were recorded in the field. Beginning with the first leaf at the tip that could be flattened and scanned (∼1 cm in length), leaves were collected in order, shoot tip to base, as a stack and placed into a Ziploc bag to which the label was affixed. A single shoot was collected per vine, and bags were placed into a cooler until scanning. A description of the number of vines (genotypes) collected for each species and hybrid, as well as the number of leaves representing different developmental stages (Sn) and leaf numbers (Ln) can be found in supporting information **(Fig. S1)**.

Leaves were arranged on a scanner (Mustek A3 1200S), in the collected order from the shoot, and next to each leaf was placed a small label indicating nodes (as measured by developmental stage) and organ identity opposite the node (“C”, cluster; “T”, tendril; “N”, no organ). The abaxial side of leaves was imaged. The file name of the image indicates the vine ID, and the appended letter indicates which image in the series the file represents for each vine. The raw scans are publically available at the following link: https://dataverse.harvard.edu/dataverse/vitis_leaves

### Landmarking

For both wild *Vitis* species and reanalyzed *V. vinifera* subsp. *vinifera* data, 17 landmarks were placed, in order, for each leaf using the ImageJ (Abramoff *et al.*, 2004) point tool. Landmarks and their order were as follows: 1) petiolar junction, 2) midvein tip, 3) left distal sinus, 4) right distal sinus, 5) left distal lobe tip, 6) right distal lobe tip, 7) left proximal sinus, 8) right proximal sinus, 9) left proximal lobe tip, 10) right proximal lobe tip, 11) left terminus petiolar vein, 12) right terminus petiolar vein, 13) branch point midvein, 14) branch point left distal vein, 15) branch point right distal vein, 16) branch point left proximal vein, 17) branch point right proximal vein. Using ggplot2 (Wickham, 2009) in R (R Core Team, 2014), graphs for landmarks from each image were visually checked for errors. If errors were detected, the landmarking was redone for those particular samples.

### Morphometric analysis and visualization

Once a quality landmarked dataset was created, a Generalized Procrustes Analysis (GPA) was undertaken using the R package shapes (Dryden, 2013). For 17 landmarks in two dimensions (x,y coordinates) for 3,292 leaves of wild *Vitis* species and 9,548 *V. vinifera* subsp. *vinifera* leaves, GPA was performed using the procGPA function, reflect=TRUE. Separate Procrustes analyses were performed for 1) all wild *Vitis* species leaves at all shoot positions and 2) *V. vinifera* subsp. *vinifera* and subsp. *sylvestris* leaves and wild *Vitis* species leaves selected for the corresponding shoot position as the domesticated grape dataset. Eigenleaves were visualized using the shapepca function and PC scores, percent variance explained by each PC, and Procrustes-adjusted coordinates were obtained from procGPA object values. To test for correlation between PCs and developmental stage or leaf number, Spearman’s rho was calculated, whereas for variability of PC values across species a Kruskal-Wallis test was used.

Linear Discriminant Analysis (LDA) on Procrustes-adjusted coordinates was performed using the lda function from the MASS package (Venables and Ripley, 2002). Species, developmental stage, and leaf number were all analyzed independent of each other. The predict function (stats package) and table function (base package) were used (dependent on MASS) to reallocate leaves (whether by species, developmental stage, or leaf number) using the linear discriminants. For the case of predicting *V. vinifera* subsp. *vinifera* developmental stage and leaf number, the wild *Vitis* species data was used as a training set to predict values for domesticated grape leaves. For the wild *Vitis* species data, for each leaf there is actual versus apparent species, developmental stage, and leaf number identities. Relative developmental stage and relative leaf number are calculated as apparent value – actual value. Mean relative developmental stage and relative leaf number values for each vine were used to determine significant deviation of species relative values from zero using a one sample, two tailed t-test. Correlation between relative values and first tendril node was analyzed using Spearman’s rho.

As previously described (Chitwood *et al.*, 2014), Germplasm Resources Information Network (GRIN) trait values, averaged on a per accession basis, for *V. vinifera* subsp. *vinifera* vines were correlated with each other and morphometric and predicted temporal data using the rcorr function from Hmisc (Harell, 2013) using Spearman’s rho and false-discovery rate controlled using the Benjami-Hochberg procedure (Benjamini and Hochberg, 1995). Hierarchical clustering for *V. vinifera* subsp. *vinifera* traits, and for averaged Procrustes-adjusted coordinates for species throughout the genus *Vitis,* was carried out using the hclust function on a distance matrix calculated from correlation performed on complete, pairwise observations and visualized using the as.phylo function from the package ape (Paradis *et al.*, 2004).

Visualization was performed in ggplot2 (Wickham, 2009) using geomj_bar, geom_jboxplot, geom_point, geom_segment, geom_tile, and stat_smooth functions, among others, and color schemes derived from colorbrewer2.org.

## Results

### Developmental stage versus leaf number

To determine the effects of developmental stage, leaf number, and evolutionary lineage (taxonomic identity) on leaf shape we scanned >3,200 leaves from wild relatives of domesticated grape held by the USDA germplasm repository in Geneva, NY. Like many living collections of perennial crops, the Geneva repository houses multiple genotypes of different *Vitis* species in common conditions. This extensive collection includes wild-collected accessions of at least 19 North American and Asian *Vitis* species as well as assorted *V. vinifera* hybrids. From the Geneva repository we sampled >270 vines representing 12 *Vitis* species, 4 *V. vinifera* hybrids, and 3 species from the related genus *Ampelopsis.* Each sampled vine represents a unique genetic accession. The number of vines sampled for each species varied widely **(Fig. S1),** but in this study we focus on 10 species for which ≥5 accessions were sampled: *Vitis riparia* (71 vines), *V. labrusca* (40 vines), *V. cinerea* (37 vines), *V. rupestris* (28 vines), *V. acerifolia* (16 vines), *V. amurensis* (16 vines), *V. vulpina* (13 vines), *V. aestivalis* (8 vines), *V. coignetiae* (5 vines), and *Vitis palmata* (5 vines). All *Vitis* leaves possess a midvein, distal and proximal veins, a petiolar vein, as well as proximal and distal lobes and sinuses, and wide variation in the width of the petiolar sinus **(Fig. 1a).** We leverage these homologous points and others to measure 17 landmarks in all leaves **(Table S1).** Later, we compare the leaves from the wild relatives of grape described above to previously published data on 1,200 varieties of domesticated grape (Chitwood *et al.*, 2014), which we describe in subsequent sections.

For each vine accession, a representative shoot was selected and the shoot position of each leaf recorded **(Fig. 1b-c).** Developmental stage is measured by counting from the youngest, first measureable leaf at the shoot tip. The time between successively initiated leaves is a plastochron, and the youngest initiated leaf primordium at the shoot apical meristem is denoted P1 (plastochron 1) to indicate this. But because we begin not with P1 (which is microns in size) but with the first measureable leaf (∼1 cm in size), we use S1 (for “stage”) to denote the youngest measured leaf at the shoot tip counting numerically upwards (S2 … Sn) towards the shoot base. Contrastingly, leaf number begins with the first initiated leaf (L1) found at the shoot base and counts numerically upward (L2 … Ln) towards the shoot tip **(Fig. 1c).** These two metrics are used to differentiate the effects of developmental stage (Sn) from leaf number (Ln) **(Fig. 1c).**

The effects of developmental stage are expected to be strongest in young leaves at the shoot tip (with low S numbers and high L numbers; **Fig. 1c**), as their shape is in flux during their expansion. Older leaves at the shoot base (with low L numbers and high S numbers) are more influenced by leaf number, because they reflect changes in mature leaf shape at successive nodes. The vast majority of the vines we sampled possess leaves corresponding to S1-S10 and L1-L10 (86% represent developmental stages up to S10 and 88% represent leaf numbers up to L10, **Fig. S1)** and we restrict our analyses to these positions **(Fig. 1c).**

It is important to note that because developmental stage (Sn) and leaf number (Ln) are counted from opposite ends of the same shoots, it is anticipated that effects associated with each will generally be inversely related. Indeed, as described subsequently, this is often the case. However, the effects of each do not perfectly mirror each other, allowing the partial discernment of developmental stage (Sn) and leaf number-specific (Ln) effects. The reason for this is that the total number of leaves per shoot varies, such that S1 does not always correspond to the same leaf number between shoots **(Fig. S2a),** and likewise L1 does not always correspond to the same developmental stage between shoots **(Fig. S2b).** Importantly, the total leaves per shoot is relatively constant between species **(Fig. S1c),** so that the overall distribution of Sn and Ln between species is not confounded with species identity. Intra-species variability in total leaves per shoot partially resolves the confounding between developmental stage and leaf number, allowing for insights into trends affecting leaf ontogeny and heteroblastic shape progression. Short of tracking the developmental progression of each leaf in the shoot of each of hundreds of vines (something that is impossible at this time), the complementary indexing using developmental stage (Sn) and leaf number (Ln) allows distinct ontogenetic and heteroblastic trends in leaf shape to be described.

### Morphospace of leaves in the genus Vitis

Landmarks are aligned (accounting for translation, rotation, and scaling) using a Generalized Procrustes Analysis (GPA). A Principal Component Analysis (PCA) is then performed to visualize the major sources of shape variance among leaves, including all species and node positions. The first four PCs explain 73.2% of shape variance among leaves from *Vitis* species, developmental stages, and leaf numbers **(Fig. 2).** More than half of the shape variance, represented by PC1 and PC2, is influenced by lobing and the width of the petiolar sinus. Low PC1 and high PC2 values readily distinguish highly dissected *Vitis* species and some members of the related genus *Ampelopsis* **(Fig. 2b).** A subset of species with diverse leaf shapes is projected onto the overall morphospace in **Fig. 2b** for clarity and to better discern the shape effects by genotype of different PCs (note: all node positions are included in this visualization). In **Fig. S3,** we project leaves found throughout all nodes in 10 *Vitis* spp. with a replication ≥5 vines **(Fig. S1b;** *V. amurensis, V. coignetiae, V. palmata, V. labrusca, V. aestivalis, V. vulpina, V. cinerea, V. rupestris, V. riparia,* and *V. acerifolia).* These 10 species exhibit unique shape differences but are more closely related in shape than the extremely lobed species (A. *brevipedunculata, A. acontifolia, V. thunbergii,* and *V. piasezkii*) depicted in **Fig. 2b.**

**Figure 2:**
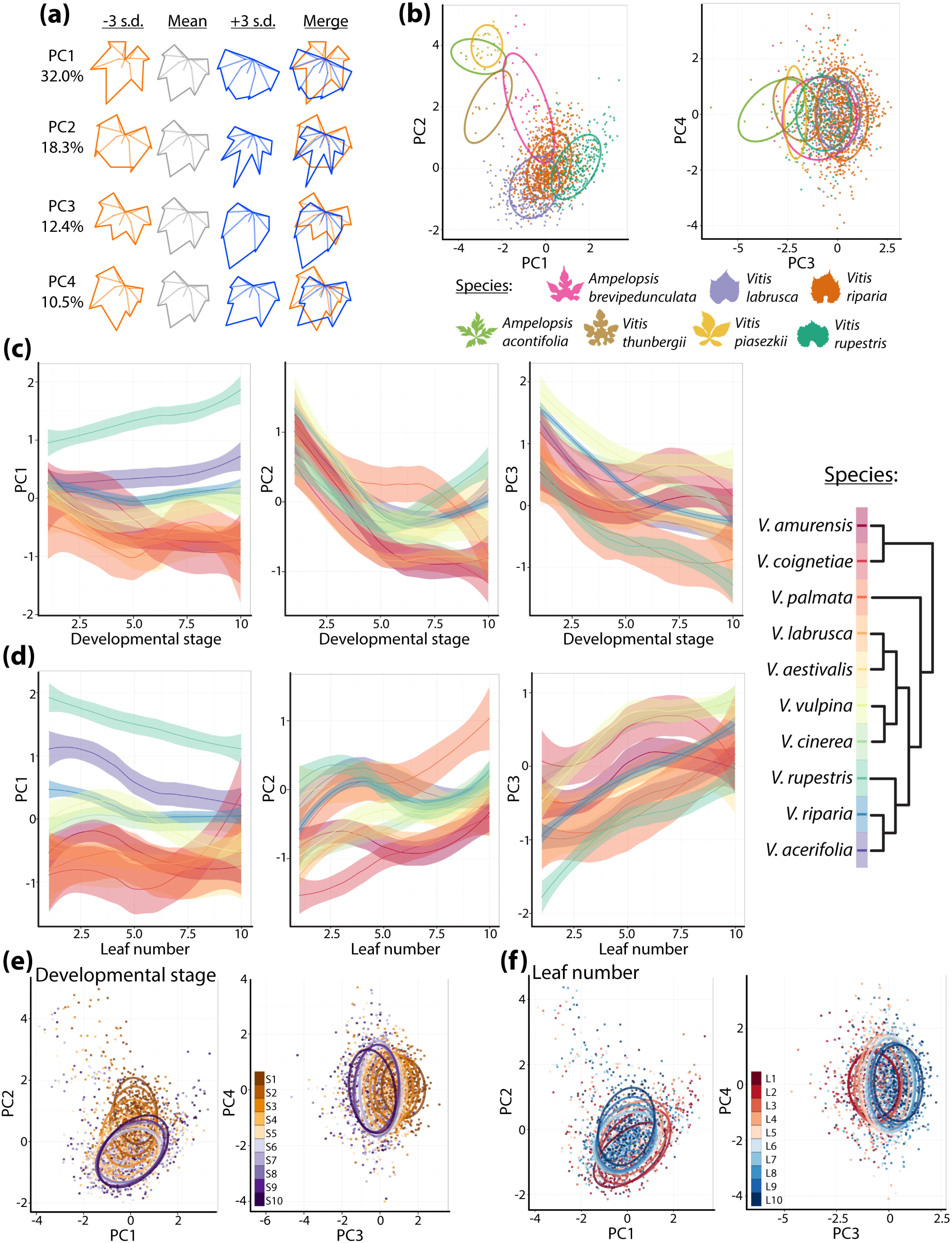
Morphospace of leaves in the genus *Vitis*. (a) “Eigenleaves” showing leaf morphs represented by Principal Components (PCs) at -/+ 3 standard deviations (s.d.) and shape variance explained by each. The PCA morphospace is calculated for all species and shoot positions, (b) Select species (indicated by color) representing morphological diversity in the dataset projected onto the morphospace. Projected data for each species includes all shoot positions. 95% confidence ellipses are drawn. Other species are projected onto the morphospace in **Fig. S3**. (c and d) Locally Weighted Scatterplot Smoothing (LOWESS) showing the relationship between PCs 1-3 and (c) developmental stage and (d) leaf number. Phylogenetic relationships between species are indicated by color. Plots of developmental stage and leaf number against each other are found in **Fig. S4.** (e) Developmental stage and (f) leaf number projected onto the morphospace representing leaves from all species.

In order to examine the variability of developmental stage and leaf number in morphospace, we visualize each PC as a locally weighted scatterplot smoothing (LOWESS) curve plotted against stage and leaf number for the 10 *Vitis* spp. for which ≥5 vines were collected **(Fig. S1b),** allowing us to statistically estimate developmental shape effects by species. Developmental stage and leaf number vary less by PC1 than by PC2 and PC3 (i.e., PC1 values are relatively developmentally-invariant compared to PC2 and PC3 values) **(Fig. 2c-d).** PC2 and PC3 values are largely indistinguishable between species across developmental stages **(Fig. 2c)** and leaf number **(Fig. 2d),** suggesting strongly conserved morphological features in developing leaves across species for these shape attributes, particularly for early developmental stages **(Fig. 2c).** Because PCs are orthogonal (i.e., uncorrelated), the fact that species, developmental stage, and leaf number correlate differentially with PCs is suggestive that each might be represented by independent shape attributes present within leaves. Re-visualizing the morphospace by developmental stage **(Fig. 2e)** and leaf number **(Fig. 2f)** demonstrates these factors mostly vary by PC2 and PC3, whereas species shape differences traverse along morphospace paths defined by PC1, PC2, and PC3 **(Fig. 2b).** Although the trajectory of shape changes in each species across developmental stage (Sn) and leaf number (Ln) are generally inversely related, this is not completely the case. In **Fig. S4,** developmental stage (Sn) and leaf number (Ln) are plotted against each other in the same plot, demonstrating their partial independence from each other.

To help qualitatively understand the different ways grape leaves differ among species and developmental contexts, we compared average shapes **(Fig. 3).** Related members of Moore’s Series Ripariae *(V. acerifolia, V. riparia,* and *V. rupestris)* (Moore, 1991; Miller *et al.*, 2013) are defined by a shallow petiolar sinus, especially *V. rupestris* **(Fig. 3a).** Outside of the Ripariae, *V. vulpina* and *V. cinerea* also exhibit shallow petiolar sinuses but to a lesser extent, while the petiolar sinus of the remaining species *(V. aestivalis, V. labrusca, V. palmata, V. coignetiae, V. amurensis*) is more acute. *Vitis palmata* exhibits especially deep distal lobing relative to other species. Each species’ leaves also vary by developmental stage **(Fig. 3b)** and leaf number **(Fig. 3c),** usually by the length of the leaf tip and the shallowness of the petiolar sinus. In the next section, we determine the extent that shape attributes varying by species, developmental stage, and leaf number are separable from each other.

**Figure 3:**
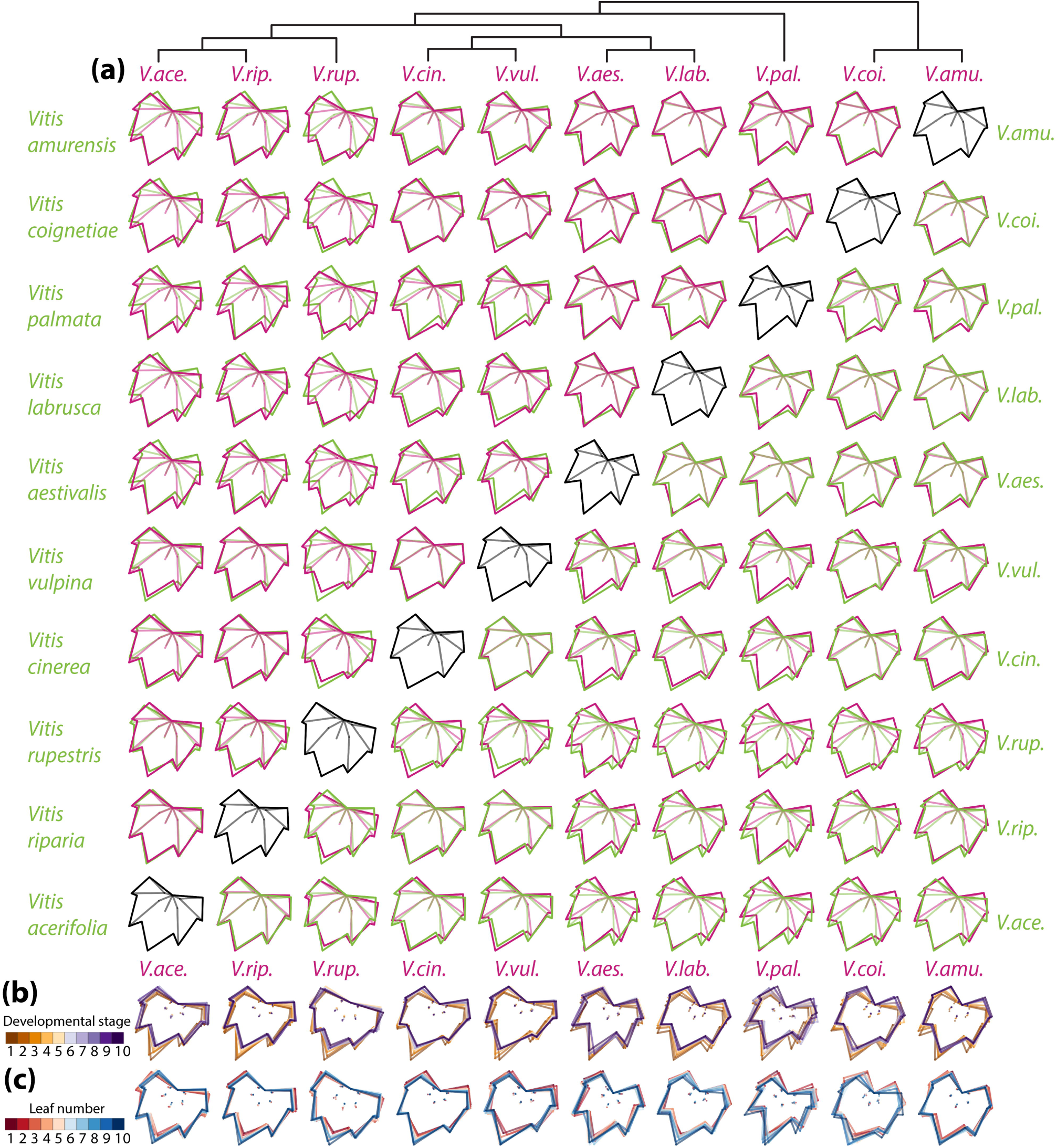
Shapes represented among *Vitis* species, developmental stage, and leaf number. (a) Outlines representing the average shape of each pairwise comparison of species, indicated by magenta and green. Phylogenetic relationships are indicated, (b and c) Comparison of average shapes across (b) developmental stages and (c) leaf number for each species indicated by color.

### Latent shapes independently predict species, developmental stage, and leaf number

We used a Linear Discriminant Analysis (LDA) to maximize the separation of leaf attributes (whether species, developmental stage, or leaf number) from each other using all measured shape information. The resulting linear discriminants can be used to predict the apparent class of a leaf and are useful for comparing actual versus apparent leaf identities as confusion matrices **(Fig. 4).**

**Figure 4:**
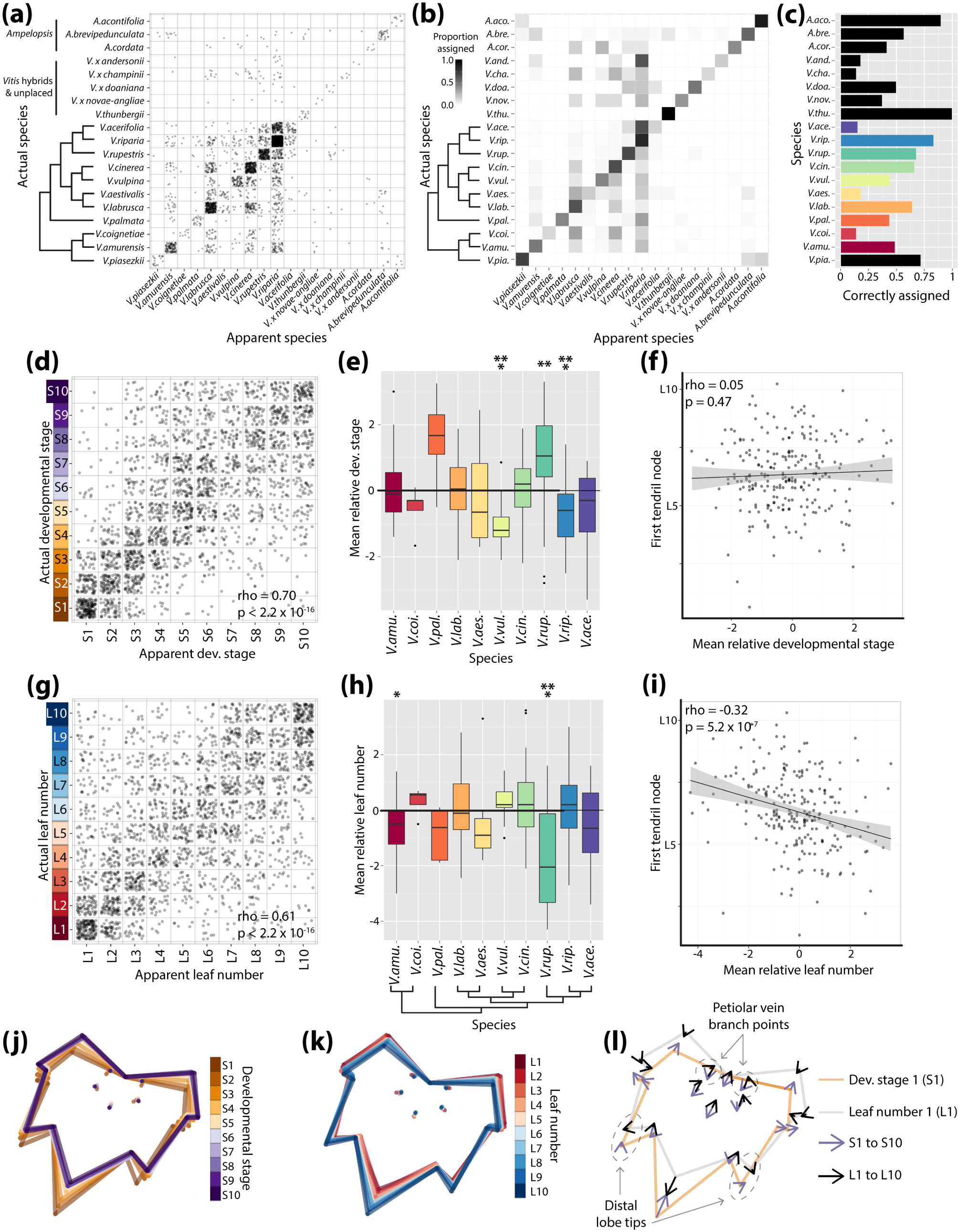
Species, developmental stage, and leaf number can be predicted independently of each other. (a) Confusion matrix resulting from prediction of species identity using a Linear Discriminant Analysis (LDA). The panel should be read left to right: for each species indicated on the left, leaves appeared to be derived from which apparent species (indicated on bottom)? Developmental stage and leaf number were not considered in the prediction, (b) Same as (a), except the apparent species assigned to each actual species have been converted into a proportion. Proportion totals equal one adding left to right, (c) The proportion of correctly assigned leaves for each species, (d) Similar to (a), actual developmental stage (left) and apparent developmental stage (bottom), predicted without species or leaf number information, (e) Relative developmental stage values (apparent - actual developmental stage) averaged for vines from each species. Significant deviations from 0 indicated by * (p<0.05), ** (p<0.01), and *** (p<0.001). (f) Correlation between mean relative developmental stage of vines and first tendril node. Fitted linear model is drawn, (g-i) same as (d-f) but for leaf number. Note the significant correlation between first tendril node and relative leaf number (i) not seen for developmental stage (f). (j and k) Latent shape attributes defining developmental stage (j) and leaf number (k) indicated with average leaf shapes. (1) Comparison of changes in latent shape attributes across developmental stage (orange, outline S1; purple, arrow to S10) and leaf number (grey, outline L1; black, arrow to L10). Note non-inverse relationship between developmental stage and leaf number changes in the distal lobe tips and petiolar vein branch points (indicated).

Linear discriminants separating species, without regard to developmental stage or leaf number, can be used to predict species identity **(Fig. 4a-c).** For most taxa, the largest proportion of predicted leaves corresponds to the taxonomic identity assigned to that species **(Fig. 4b,** the diagonal indicates the proportion of correctly assigned leaves). Incorrectly assigned leaves are most often confused for other species that are phylogenetically related to the assigned species. For example, a majority of *V. acerifolia* leaves are confused for its close relative *V. riparia*; similarly, the largest group of *V. aestivalis* leaves are confused for its relative *V. labrusca.* In contrast, the Asian *V. coigenetiae* leaves are confused for more distant relatives native to North America, including *V. labrusca* and *V. cinerea* and sometimes even *V. riparia,* perhaps indicating convergent evolution, germplasm misidentification, or segregating leaf shape differences among these accessions. Interestingly, *Vitis xandersonii,* described as a hybrid of *V. coignetiae* and either *V. vulpina* or *V. riparia,* is strongly mistaken for *V. riparia,* providing circumstantial evidence of parentage. As has been suggested by ampelographers and taxonomists (Moore, 1991; Galet, 1952), our results demonstrate that leaf shape is often sufficient to identify species, even in the absence of developmental information. Instances where leaves of one species are assigned to another offer a valuable opportunity to develop hypotheses about phylogenetic history, hybridization, and convergent evolution that can be tested with additional evolutionary and ecological analyses.

Reciprocally, we wondered whether developmental stage and leaf number, regardless of species identity, might be similarly informative about developmental context. Indeed, linear discriminants trained on developmental stage or leaf number can predict the developmental context of a leaf for either measure **(Fig. 4d-4i)**. Developmental stage can be predicted with a Spearman’s correlation coefficient of rho = 0.70 **(Fig. 4d)**, and leaf number with rho = 0.61 **(Fig. 4g)**. Prediction of both developmental stage and leaf number is more accurate at the beginning of their respective series, where the effects of each are anticipated to be the strongest **(Fig. 1c)**. Based on these observations we conclude that developmental context, measured by either developmental stage or leaf number, can be predicted independently of genotypic information. These results suggest that the allometric changes in leaf shape during ontogeny and the changes in shape due to heteroblastic development are broadly conserved across the genus *Vitis.*

The ability to predict developmental context separate from species identity provides a method to quantify differences between species attributable to changes in developmental timing, also known as heterochrony. For each leaf, we calculate relative developmental stage and relative leaf number as the apparent value minus the actual value (e.g., an S5 leaf predicted to be S7 would have a relative stage value of +S2, and an L4 leaf predicted to be L2 would have a relative value of –L2). Relative values indicate, for a given leaf, how many nodes ahead or behind (developmentally speaking) that leaf appears to be. Averaging the relative developmental stage and leaf number values for each vine, we can detect species that are precociously ahead or lagging behind from expected developmental stage **(Fig. 4e)** and leaf number values **(Fig. 4h)**. Relative developmental stage and relative leaf number are inversely related to each other (compare **Fig. 4e** with **Fig. 4h**) but not completely so, consistent with the Sn and Ln numbers showing inversely-related but unique trajectories through morphospace **(Fig. S4)**. Relative developmental stage and leaf number provide contrasting insights into the temporal development of leaves. For example, the mean relative developmental stage of *V. rupestris* vines is about +1S ahead of their actual developmental stage **(Fig. 4e)** and about -2L behind actual leaf number (**Fig. 4h**). This accelerated development (Sn) and protracted display of juvenile leaf types typically found at the base of the shoot (Ln) contributes towards the unique shape of *V. rupestris* leaves compared to other *Vitis* spp. **(Fig. 3)**. *Vitis rupestris* leaves **(Fig. 3a)**, older leaves (high Sn values, **Fig. 3b)**, and juvenile leaves (low Ln values, **Fig. 3c**) all share a characteristic wideness conferred by a diminished leaf tip.

Within *Vitis,* the identity of inflorescences, which appear opposite leaves, transform from clusters at the shoot base to tendrils at the shoot tip (Srinivasan and Mullins, 1981; Gerrath, 1988; Gerrath, 1993; Boss and Thomas, 2002) **(Fig. 1c)**. Both changes in leaf shape at successive nodes and the transformation of inflorescence identity indicate temporal changes in the development of the shoot apical meristem, known as heteroblasty. This transition is not the same as flowering time, as both clusters and tendrils are inflorescences, and moreover these organ primordia are patterned the previous year (Carmona et al., 2008). Therefore, the cluster-to-tendril transformation serves as a discrete, binary indication of the heteroblastic transition, i.e., the temporal development of the meristem. If heteroblasty regulates both the latent shapes predicting leaf number (Ln) and the cluster-to-tendril transition, we would assume they would be correlated. Further, such a correlation should not be observed between developmental stage (Sn) and the cluster-to-tendril transition, as leaf ontogeny (developmental stage) is not related to heteroblasty (cluster-to-tendril transition).

Indeed, there is a significant negative correlation between relative leaf number and the first node a tendril is observed: that is, in vines with precocious, adult leaves (i.e., higher relative leaf numbers, typical of leaves found more towards the shoot tip), tendrils appear at earlier nodes closer to the base of the shoot, linking two different measures of premature heteroblastic change (**Fig. 4i)**. Moreover, this correlation is not observed for relative developmental stage **(Fig. 4f)**. Taken together, these results indicate that latent shapes predictive of developmental stage and leaf number are functionally distinct, reflecting the developmental progression of leaves (Sn) and their heteroblastic transitions across successive nodes of the shoot (Ln) separately.

Average leaf shapes indicate that developmental stage and leaf number vary by the prominence of the leaf tip and the shallowness of the petiolar sinus **(Fig. 4j-l**). The shape changes for these two factors are inversely related, but obviously distinguishable from each other, given their differential correlation with tendril position, i.e. heteroblasty **(Fig. 4f,4i**). Upon closer inspection, the distal lobe tips and the branch point of the petiolar vein distinguish shape changes attributable to developmental stage and leaf number (**Fig. 4l**), isolating the shape attributes unique to these functionally distinct processes.

### Morphological differences between domesticated grape and wild relatives

Previously, we measured >9,500 leaves from *V. vinifera* subsp. *Vinifera* (domesticated grape) and its wild progenitor *V. vinifera* subsp. *sylvestris.* These leaves, collected from the USDA germplasm repository in Winters, CA, represent over 2,300 vines and 1,200 varieties (Chitwood *et al.*, 2014). These leaves were sampled to measure purely genetic effects and minimize the influence of development by collecting four successive leaves from the midpoint of the shoot. That species identity can be predicted independently from developmental context in wild *Vitis* spp. **(Fig. 4a-c**) with disparate leaf shapes lends credence to the assumption that developmental context can be ignored, which is almost always invoked when mapping traits for quantitative genetic purposes or taxonomically defining species.

We wanted to compare leaves from domesticated grape and its wild progenitor to the other *Vitis* species described above. To do so, we restricted our analysis *in silico* to leaves at the same shoot positions we had originally collected in domesticated grape (the four leaves closest to the midpoint of the shoot). In the combined morphospace, PC1 and PC2 describe nearly 60% of shape variance **(Fig. 5)**. Similar to the wild *Vitis* species-only morphospace **(Fig. 2a)**, lobing and the petiolar sinus define the first PCs. Petiolar veins in domesticated grape varieties can be so angled they cross each other, and because of this the closure of the petiolar sinus represented by high PC1 values is particularly strong **(Fig. 5a)**. Leaf dissection defined by high PC1 and high PC2 values clearly delineates less lobed wild *Vitis* species from more acutely lobed domesticated grape varieties **(Fig. 5b)**. Highly dissected species such as *Ampelopsis acontifolia, V. piasezkii, V. thunbergii,* and *V. vinifera* var. Ciotat form a distinct morphological grouping.

**Figure 5:**
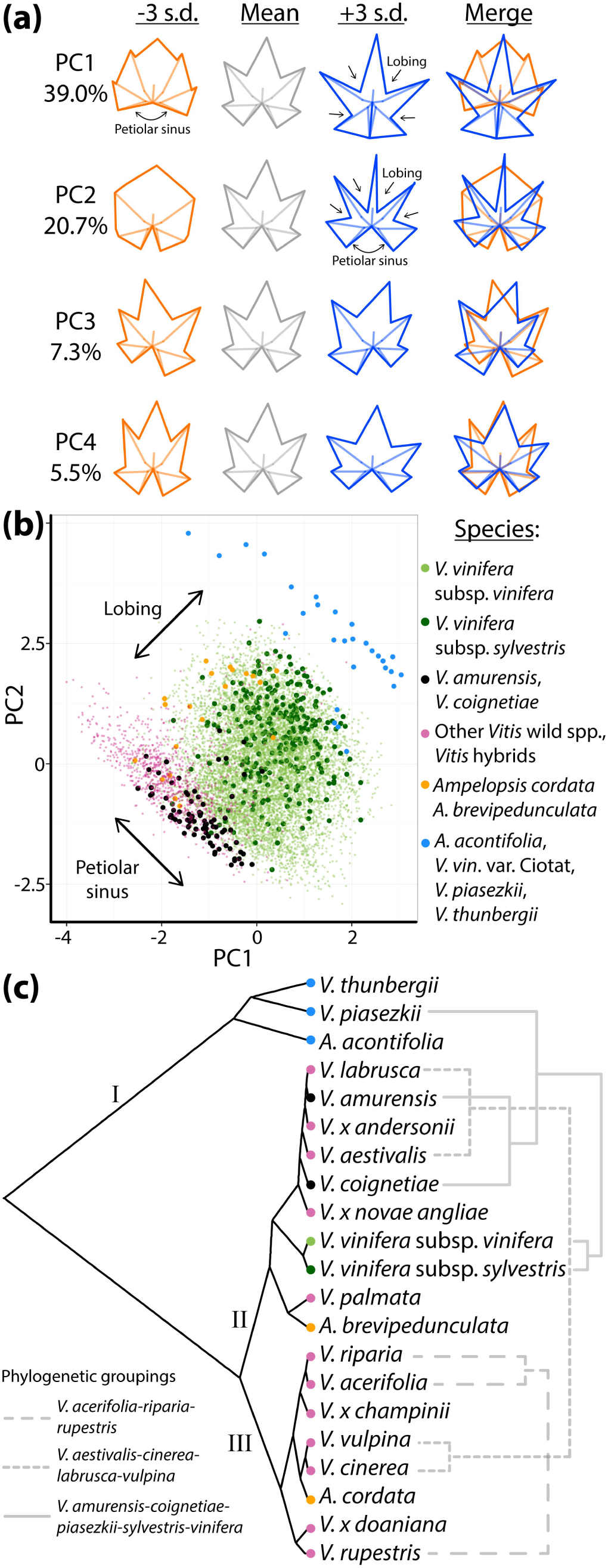
Morphospace of wild *Vitis* species and domesticated grape. (a) “Eigenleaves” showing leaf morphs represented by PCs at -/+ 3 s.d. and shape variance explained by each. Lobing and petiolar sinus variation indicated, (b) Comparison of wild *Vitis* species leaf shape (red, black) with *V. vinifera* subsp. *vinifera* and *V. vinifera* subsp. *sylvestris* (green, dark green, respectively), and highly-lobed species (blue). Strong shape variation in lobing and petiolar sinus depth indicated, (c) Hierarchical clustering based on morphology. Phylogenetic groupings indicated in light gray. Groups mentioned in text indicated with Roman numerals.

While there are clear patterns in leaf shape related to development and species identity, correspondence between the morphospace of wild and domesticated *Vitis* species and evolutionary relationships among *Vitis* species is complex. Some, but not all, well-known phylogenetic relationships are reflected in leaf morphology (**Fig. 5c**) (Miller *et al.*, 2013; Zecca *et al.*, 2012). Notably, clustering averaged shapes from different genotypes reveals that *V. vinifera* subsp. *vinifera* is morphologically sister to *V. vinifera* subsp. *sylvestris,* the wild progenitor of the domesticated grape **(Fig. 5c,** Group II) (Zecca *et al.*, 2012; Myles *et al.*, 2011). Also within Group II are two Asian species, *V. amurensis* and *V. coignetiae,* which together with *V. vinifera* subsp. *vinifera* and *V. vinifera* subsp. *sylvestris* represent members of a Eurasian clade. Within Group III are found members of Series Ripariae (*V*. *acerifolia, V. riparia, V. rupestris).* Leaf shapes of *Vitis* hybrids cluster based on parental lineages, with *V. xchampinii* and *V. xdoaniana* clustering in Group III presumably because of their *V. rupestris* and *V. acerifolia* heritages (respectively) and *V. xandersonii* and *V. xnovae-angliae* clustering closer with their *V. coignetiae* and *V. labrusca* parents (respectively). These observations suggest similarities in leaf shape may reflect recent evolutionary events, domestication, or contemporary interspecific gene flow.

At broader levels of phylogenetic scale, however, patterns of leaf shape similarity do not appear to correspond with known evolutionary relationships. For example, together with *V. vinifera* subsp. *vinifera, V. vinifera* subsp. *sylvestris,* and two Asian species found Group II (discussed above) are other North American species *(V. labrusca, V. palmata*), several hybrids, and *Ampelopsis brevipedunculata,* taxa that are not known to be closely related to one another. A previously identified clade based on molecular data *(V. aestivalis, V. cinerea, V. labrusca,* and *V. vulpina)* here is divided between Groups II and III. Although *Vitis* is well known to be a monophyletic genus (Wen *et al.*, 2007; 2013), based on leaf shape three species of *Ampelopsis* cluster in three different groups, together with *Vitis* species. These data suggest that while leaf shape may bear signatures of recent evolutionary events, leaf shape does not appear to track with phylogeny at larger scales; distantly related species, even from distinct genera, resemble each other in leaf shape through evolutionary convergence.

### Changes in developmental timing underlie morphological diversity in domesticated grape leaves

Even though leaves from equivalent positions in the shoot clearly separate species and varieties by genetic effects **(Fig. 5),** such effects may still be developmental in nature, an example of heterochrony. If changes in the relative timing of developmental stage (Sn) or leaf number (Ln) have occurred, they would contribute to morphological differences between species, as described above for wild *Vitis* spp. **(Fig. 4e, 4h)**. The conserved latent shapes defining developmental stage and leaf number in morphologically disparate *Vitis* species can be used predictively to detect such changes in timing.

We used wild species leaves, from S1-S10 and L1-L10 (where developmental stage and leaf number effects are the strongest, respectively) as a training set, to predict the stage (Sn) and leaf number (Ln) values of domesticated grape leaves **(Fig. 6).** A caveat of this approach is that the two datasets are collected from different environments and years. When using the Geneva, NY 2013 dataset to predict developmental timing in the Winters, CA 2011 dataset, an assumption is that environment affects leaf morphology independently from developmental timing. Additionally, the wild species training set is influenced by strong morphological trends at the tip and base of the shoot, whereas the domesticated dataset is collected from the middle of the shoot where these effects are weaker. Nonetheless, developmental timing can be predicted independently from genotype in the middle of the shoot as well, even if the effects are weaker than at the ends of the shoot (see **Fig. 4d, 4g).**

**Figure 6:**
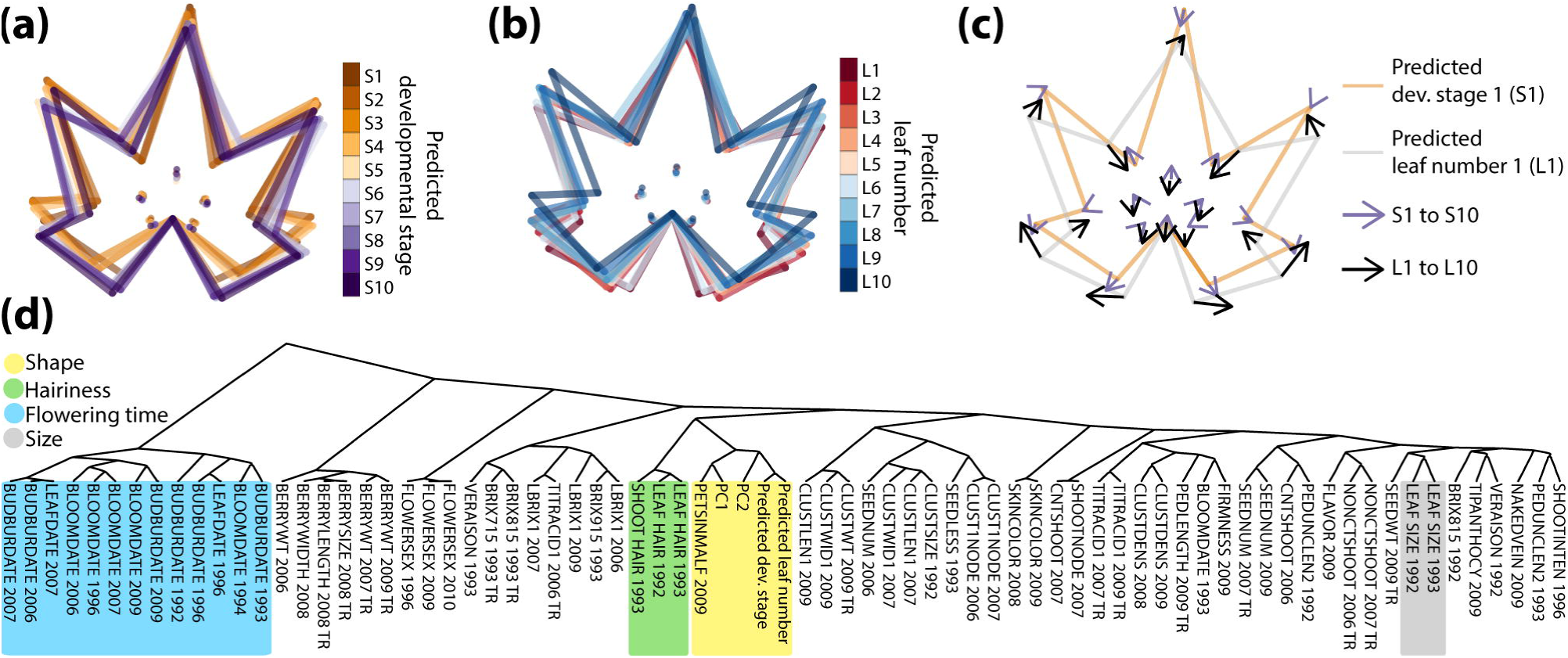
Evolution in developmental timing underlies leaf shape diversity in domesticated grape. (a and b) Latent shape attributes defining developmental stage (a) and leaf number (b) in domesticated grape indicated with average leaf shapes, (c) Comparison of changes in latent shape attributes across predicted developmental stage (orange, outline S1; purple, arrow to S10) and predicted leaf number (grey, outline L1; black, arrow to L10). (d) Hierarchical clustering of measured traits in grape. Morphological and predicted temporal traits (yellow), hirsuteness (green), reproductive transitions (blue), and overall leaf size (grey) indicated.

When comparing leaf averages across predicted developmental stages **(Fig. 6a)** and leaf numbers **(Fig. 6b),** it is apparent that variation in petiolar sinus depth is the major predictor of developmental timing in domesticated grape leaves **(Fig. 6c).** Variation in the petiolar sinus is a recurrent theme in morphology associated with both developmental stage and leaf number **(Figs. 3b-c; 4j-l).**

Previously, numerous other traits, from timing of bud burst and flowering to berry ripening and titratable acids and sugar content, had been measured on the domesticated grapevines described above (Chitwood *et al.*, 2014). We were curious about the correlational context of overall leaf shape (represented by PCs) and developmental timing (predicted developmental stage and leaf number) to these other traits. The predicted developmental stages and leaf numbers of domesticated grape leaves are significantly correlated with PC1 and PC2 **(Table S2; Fig. 6d),** which together explain near 60% of total shape variance for measured *Vitis* genotypes **(Fig. 5a).** From this, we conclude that latent shape attributes modulating developmental stage and leaf number potentially explain large amounts of shape variance in domesticated grape, indicating changes in developmental timing (heterochrony) may underlie the morphological evolution of domesticated grape varieties. Additionally one trait, “PETSINMALF 2009”, is a qualitative, 0-8 rating of petiolar sinus depth, and is tightly correlated to PC1, PC2, and predicted developmental stage and leaf number, confirming that our quantitative measures reflect intuitive perceptions of leaf shape. **(Fig. 6d; Table S2).**

Leaf morphology traits, including predicted developmental timing, are largely separate from bud burst and bloom dates (“BUDBURST DATE”, “BLOOMDATE”, “LEAFDATE”), indicating that these timing features are separate from the overall transition to reproductive development in grapevine **(Fig. 6d).** Further, leaf shape is distinct from overall size of leaves (“LEAF SIZE”), demonstrating that grape leaf shape is not merely a developmental constraint correlated with overall size (i.e., allometry) **(Fig. 6d).** One group of traits significantly correlated and clustering with leaf shape is hirsuteness, both in leaves and the shoot (“LEAF HAIR”, “SHOOT HAIR”), which we had observed previously in domesticated grape (Chitwood *et al.*, 2014). Leaf hirsuteness varies by both developmental stage, as trichomes become less dense as the leaf expands, and also by leaf number. It is interesting to think about how thermoregulation, disease resistance, or other trichome mediated functionality may vary as a developmental constraint as developmental timing changed over the evolution and domestication of grapevines.

## Discussion

Many descriptions of complex traits, from faces to voices, and even the evolution of cultural products, such as violins (Zhand *et al.*, 1997; Kuhn *et al.*, 2000; Claes *et al.*, 2014; Chitwood, 2014), potentially stand to benefit from isolating features uniquely regulated by distinct pathways. Like a face, leaf shape is modulated in complicated ways by genetics, development, and the environment. The latent shapes we describe here, describing developmental stage and leaf number, embedded within the overall shape of a leaf, and conserved across the genus *Vitis,* fit the definition of a “cryptotype”. Recently, we had suggested the term “cryptotype,” borrowed from linguistics (which uses both “phenotype” and “cryptotype” parallel to the biological definitions) (Whorf, 1945), to describe latent, combinatorial features (e.g., 17 homologous landmarks) within a shared multivariate space that vary independently from each other (Chitwood and Topp, 2015). Cryptotypes are not unreal or even abstract; they are simply a set of features, that when combined, best discriminate one biological process (e.g., evolutionary history) from others (e.g., developmental stage). The shapes that arise through this combination of features are latent only because they are not immediately recognizable when each of their component landmarks is considered alone; the effects of genetics and development affect the entirety of the leaf. The multivariate cryptotypes of genotype, developmental stage, and leaf number stand in contrast to traditional univariate phenotypes, such as height, biomass, or leaf length, width, or area.

Defining such combinative, latent shapes is critical to understand the mechanisms by which morphological diversity manifests in plants. The unequal expansion of leaves (allometry) and the different types of leaves displayed at nodes (resulting from heteroblasty) have been studied for centuries (Hales, 1727; Goethe, 1817; Goebel, 1900; Goethe (trans.), 1952), but the inability to distinguish the effects at a morphological level, within the shared morphology of single leaves where these processes act, has led to confusion. Early hypotheses that shade prolongs juvenility through a heteroblastic mechanism were later refuted by carefully studying the morphology of leaf primordia (Jones, 1995). Similar, the degree that changes in the development of leaves or the heteroblastic series influences evolutionary changes—and plasticity—remains an open question.

Recent studies have uncovered a transcriptomic basis underlying leaf ontogeny (Efroni *et al.*, 2008). Similarly, dramatic transitions to reproductive fates reflecting the temporal development of meristems have been described (Park *et al.*, 2012). An understanding of molecular clocks, regulating phase change through small RNAs (Chuck *et al.*, 2007; Wu *et al.*, 2009), and their signals, such as sugar (Yang *et al.*, 2013; Yu *et al.*, 2013), has come into focus. Vegetative, heteroblastic changes in leaf shape have an intimate relationship with reproductive transitions, as demonstrated by recent work linking natural variation in a floral repressor, FLC, with changes in leaf morphology between species (Cartolano et al., 2015). Of the many gene regulatory networks regulating leaf shape, specific pathways have been identified mediating heteroblastic shape changes (Rubio-Somoza *et al.*, 2014). Similar molecular descriptions of leaf development exist (Ichihashi *et al.*, 2014) and the crosstalk between these two processes is large. Strangely, even though molecular correlates of leaf development and heteroblasty have been uncovered, a phenotypic basis for the morphological effects of temporal patterning remains obscure, except in a qualitative sense.

Decomposing complex morphologies into latent shapes regulated by distinct pathways is required to precisely describe the mechanisms that, together, produce the composite shape of a single leaf. By isolating latent shapes that are regulated by evolution and development, we can focus on those attributes of the leaf most relevant for producing the diversity of leaf shapes observed in nature, and consider the functional contributions of leaf morphology to assessments of biodiversity, plant responses to global climate change, and the genetic improvement of crop species.

## Acknowledgements

This work was funded by research stipends awarded to DHC, LLK, and JPL through the Grape Research Coordination Network, DBI0741876. DHC, LLK, AJM, JPL designed the research; DHC contributed new analytic/computational/etc. tools; DHC, LLK, RO, SC, MG, CK analyzed data; DHC, LLK, AJM, JPL wrote the paper.

## Supplemental Data

**Figure S1:** Numbers of species, vines, leaves, and shoot positions sampled.

**Figure S2:** Developmental stage and leaf number are partially confounded.

**Figure S3:** Projection of species with ≥ 5 vines onto the morphospace.

**Figure S4:** Comparison of leaf shape changes due to developmental stage (Sn) and leaf number (Ln) for different species.

**Table S1:** Procrustes-aligned coordinates.

**Table S2:** Correlation between traits.

